## Supplementary Information for "3D Cell Aggregates Amplify Diffusion Signals"

Spheroids were transferred to a 96-well flat bottom ultra-low attachment plate (Corning) and medium was changed to 75 or 100 $\mu$L (8 replicates per volume) HepaRG medium with 11.1 mM glucose and 870 nM insulin. Medium samples were taken after 1, 5, and 10 min and HepaRG medium with 11 mM glucose and 870 nM insulin was sampled as 0 min control. For samples where starting volume was 100 $\mu$L, a medium sample was also taken after 4 h. For samples with starting volume of 75 $\mu$L, all medium was removed after 10 min sampling and 100 $\mu$L new medium was added. From these incubations, medium was sampled after 19 h. The measured data is provided in the following tables.

**Supplementary Table S1**: Measured glucose concentration of medium before adding the spheroids

| | | | | | | | |
|---|---|---|---|---|---|---|---|
| 11.34 | 11.15 | 10.49 | 11.79 | 11.21 | 10.95 | 11.26 | 10.76 |

**Supplementary Table S2**: Measured Glucose concentration at time points 1, 5, 10 min, and 4h in 100 $\mu$L incubations.

| Replicate number | 1 | 2 | 3 | 4 | 5 | 6 | 7 | 8 |
|---|---|---|---|---|---|---|---|---|
| **1min** | 9.94 | 11.05 | 10.95 | 10.85 | 11.32 | 10.91 | 10.74 | 10.33 |
| **5min** | 12.09 | 11.03 | 11.35 | 11.46 | 10.92 | 10.92 | 11.31 | 10.43 |
| **10min** | 10.45 | 10.09 | 10.15 | 9.82 | 9.89 | 9.53 | 10.62 | 9.91 |
| **4h** | 10.24 | 10.00 | NV [a] | 9.72 | 9.77 | 9.75 | 9.25 | 8.43 |

[a] No value.

**Supplementary Table S3**: Measured Glucose concentration (GC) at times 1, 5, 10 min, and 19h in 75 $\mu$L incubations.

| Replicate number | 1 | 2 | 3 | 4 | 5 | 6 | 7 | 8 |
|---|---|---|---|---|---|---|---|---|
| **1min** | 9.96 | 11.00 | 9.89 | 10.64 | 10.58 | 9.67 | 10.96 | 10.93 |
| **5min** | 12.12 | 11.19 | 10.68 | 11.27 | 11.51 | 11.23 | 11.07 | 9.83 |
| **10min** | 10.71 | 10.69 | 11.68 | 11.15 | 11.04 | 10.97 | 11.31 | 10.61 |
| **19h** | 6.50 | 5.41 | 4.72 | 4.84 | 5.19 | 5.31 | 4.87 | 5.40 |

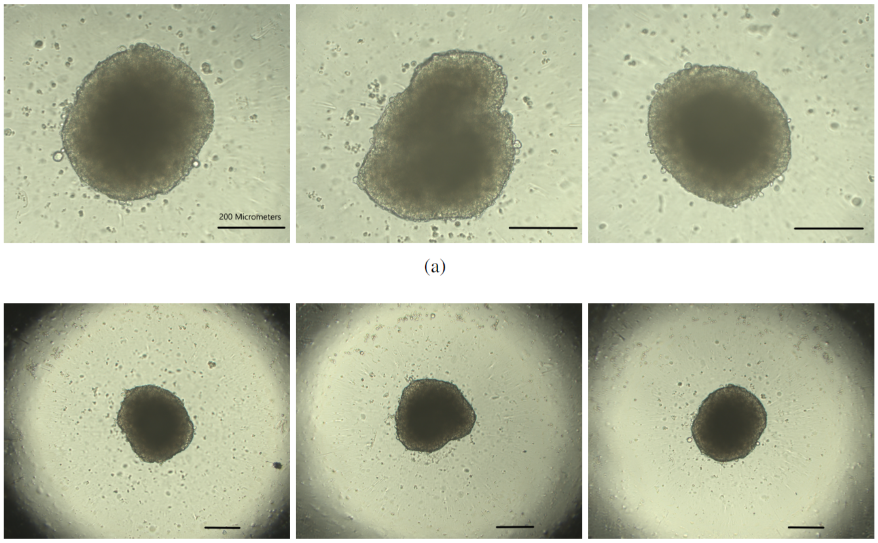


(a)

Supplementary Figure S1: (a) Microscopic images of three liver spheroids at 10X magnification (scale bar is 200 $\mu$m). (b) Microscopic images of three spheroids at 4X magnification (scale bar is 200 $\mu$m).

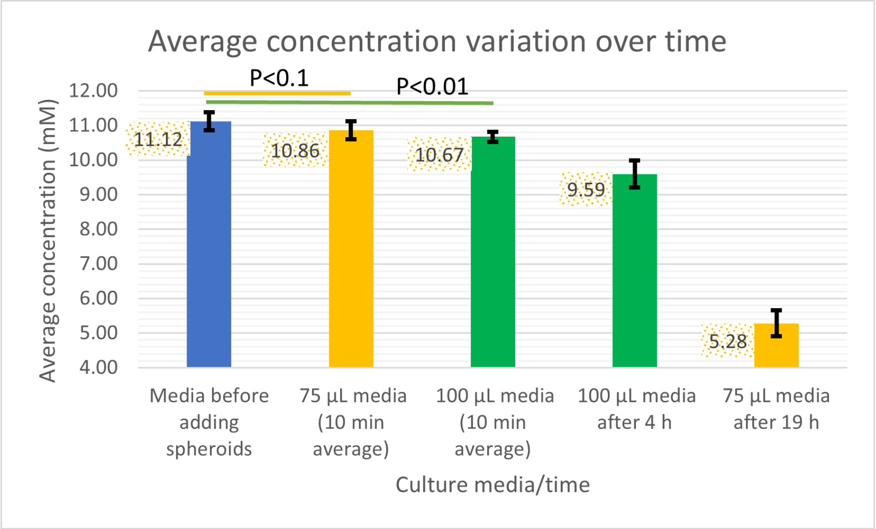


**Supplementary Figure S2:** Mean GC in medium before adding spheroids, mean 10 min time-averaged GC for the 100 and 75 $\mu$L incubations, mean GC at 4 hours (for 100 $\mu$L), mean GC at 19 hours (for 75 $\mu$L).

**Appendix S1**: *Proof of Proposition 1*: During a very short time interval $\Delta t$, the molecules within the left side region $x \in [-3\sigma_o, 0]$ (where $\sigma_o = \sqrt{2D\Delta t}$) have a realistic chance to pass across the boundary to the right. Within a small region with length $\mathrm{d}x$ at any point $x \in [-3\sigma_o, 0]$, we have $c_o(x)\mathrm{d}x$ molecules that may cross the boundary with probability $\int_{-x}^{+\infty} \frac{1}{\sqrt{2\pi}\sigma_o} \exp\left(-\frac{u^2}{2\sigma_o^2}\right) \mathrm{d}u$. Therefore, the total number of molecules crossing the boundary from the left side over $\Delta t$ seconds is $\epsilon_s \int_{-3\sigma_o}^{0} \int_{-x}^{+\infty} \frac{c_o}{\sqrt{2\pi}\sigma_o} \exp\left(-\frac{u^2}{2\sigma_o^2}\right) \mathrm{d}u\mathrm{d}x$. This can be rewritten as $\epsilon_s \int_0^{3\sigma_o} \int_{|x|}^{+\infty} \frac{c_o}{\sqrt{2\pi}\sigma_o} \exp(-\frac{u^2}{2\sigma_o^2})\mathrm{d}u\mathrm{d}x$.

Similarly, we can derive an expression for the number of molecules that move from the right to the left side of the boundary as $\epsilon_s \int_0^{3\sigma_s} \int_{|x|}^{+\infty} \frac{c_s}{\sqrt{2\pi}\sigma_s} \exp(-\frac{u^2}{2\sigma_s^2})\mathrm{d}u\mathrm{d}x$. On the other side, the molecule flux at the boundary is given by (3) in one dimension, i.e., $D\frac{dc_o}{dx}$. During a very short time interval $\Delta t$, this flux leads to the passage of $D\frac{\mathrm{d}c_o}{\mathrm{d}x}\Delta t$ molecules across the boundary. Thus, we can write

$$\epsilon_s \int_0^{3\sigma_o} \int_{|x|}^{+\infty} \frac{c_o}{\sqrt{2\pi}\sigma_o} \exp\left(-\frac{u^2}{2\sigma_o^2}\right) \mathrm{d}u\mathrm{d}x - \epsilon_s \int_0^{3\sigma_s} \int_{|x|}^{+\infty} \frac{c_s}{\sqrt{2\pi}\sigma_s} \exp\left(-\frac{u^2}{2\sigma_s^2}\right) \mathrm{d}u\mathrm{d}x = D\frac{\mathrm{d}c_o}{\mathrm{d}x}\Delta t. \tag{S1}$$

By making the substitution $u = \frac{\sigma_s}{\sigma_o}u'$ in the second term on left side of (S1) and defining $F(x) = \int_{|x|}^{+\infty} \frac{1}{\sqrt{2\pi}\sigma_o} \exp\left(-\frac{u^2}{2\sigma_o^2}\right) \mathrm{d}u$, we obtain

$$\epsilon_s \int_0^{3\sigma_o} c_o(x)F(x)\mathrm{d}x - \epsilon_s \int_0^{3\sigma_s} c_s(x)F\left(\frac{\sigma_o}{\sigma_s}x\right) \mathrm{d}x = D\frac{\mathrm{d}c_o}{\mathrm{d}x}\Delta t. \tag{S2}$$

By making the substitution $x' = \frac{\sigma_o}{\sigma_s}x$, we get

$$\int_0^{3\sigma_o} c_o(x)F(x)\mathrm{d}x - \int_0^{3\sigma_o} c_s\left(\frac{\sigma_s}{\sigma_o}x'\right)\frac{\sigma_s}{\sigma_o}F(x')\mathrm{d}x' = \frac{D}{\epsilon_s}\frac{\mathrm{d}c_o}{\mathrm{d}x}\Delta t. \tag{S3}$$

For a very short $\Delta t$, as $\sigma_0$ and $\sigma_s$ approach zero, we can approximate $c_o(x)$ and $c_s(x)$, $x \in \{0, 3\sigma_0\}$ as constant values representing the border concentrations at the right and left sides of boundary denoted by $c_o(0^-)$ and $c_s(0^+)$ respectively within the integral bounds of $[0, 3\sigma_0]$ and $[0, 3\sigma_s]$. We note that as $\Delta t$ approaches 0, $\sigma_0$ and $\sigma_s$ tend to 0 which enables arbitrarily approaching the boundary from both sides, rather than restricting the proof to a specific time. Therefore, (S3) leads to

$$\left(c_o(0^-) - c_s(0^+)\frac{\sigma_s}{\sigma_o}\right) \int_0^{3\sigma_o} F(x)\mathrm{d}x = \frac{D}{\epsilon_s}\frac{\mathrm{d}c_o}{\mathrm{d}x}\Delta t. \tag{S4}$$

Note that a change of variable simplifies $F(x)$ to $\int_{|x|/\sigma_o}^{+\infty} \frac{1}{\sqrt{2\pi}} \exp(-\frac{u^2}{2})du = Q(|x|/\sigma_o)$ where $Q(\cdot)$ is the Q-function. Furthermore, by one additional variable change, we obtain

$$\int_0^{3\sigma_o} F(x)dx = \int_0^{3\sigma_o} Q(|x|/\sigma_o)dx = \sigma_o \int_0^{3} Q(x)\mathrm{d}x \simeq 0.4\sigma_o.$$

Therefore, (S4) is written as

$$\left( c_o(0^-) - c_s(0^+)\frac{\sigma_s}{\sigma_o} \right) 0.4\sigma_o = \frac{D}{\epsilon_s}\frac{\mathrm{d}c_o}{\mathrm{d}x}\Delta t. \tag{S5}$$

Considering $D\Delta t = \sigma_o^2/2$, we obtain

$$\left( c_o(0^-) - c_s(0^+)\frac{\sigma_s}{\sigma_o} \right) 0.8\epsilon_s = \frac{\mathrm{d}c_o}{\mathrm{d}x} \times \sqrt{2D\Delta t}. \tag{S6}$$

Obviously, as $\Delta t \to 0$, the right side goes to zero, which implies that $\sigma_o c_o(0^-) = \sigma_s c_s(0^+)$ or equivalently $k = \frac{\sigma_o}{\sigma_s} = \sqrt{\frac{D}{D_{\text{eff}}}}$. This completes the proof.
